# Regional random mutagenesis driven by multiple sgRNAs and diverse on-target genome editing events to identify functionally important elements in non-coding regions

**DOI:** 10.1101/2023.09.12.557308

**Authors:** Kento Morimoto, Hayate Suzuki, Akihiro Kuno, Yoko Daitoku, Yoko Tanimoto, Kanako Kato, Kazuya Murata, Fumihiro Sugiyama, Seiya Mizuno

**Author notes:** To whom correspondence should be addressed., Correspondence may also be addressed to Seiya Mizuno.

## Abstract

Functional regions that regulate biological phenomena are interspersed throughout eukaryotic genomes. The most definitive approach for identifying such regions is to confirm the phenotype of cells or organisms in which specific regions have been mutated or removed from the genome. This approach is invaluable for the functional analysis of genes with a defined functional element, the protein-coding sequence. In contrast, no functional analysis platforms have been established for the study of cis-elements or microRNA cluster regions consisting of multiple microRNAs with functional overlap. Whole-genome mutagenesis approaches, such as via *N*-ethyl-*N*-nitrosourea (ENU) and gene trapping, have greatly contributed to elucidating the function of coding genes. These methods almost never induce deletions of genomic regions or multiple mutations within a narrow region. In other words, cis-elements and microRNA clusters cannot be effectively targeted in such a manner. Herein, we established a novel region-specific random mutagenesis method named <u>C</u>RISPR-& <u>T</u>ransposase-based <u>R</u>egiona<u>L</u> Mutagenesis (CTRL-Mutagenesis). We demonstrated that CTRL-mutagenesis randomly induces diverse mutations within target regions in murine embryonic stem cells. Comparative analysis of mutants harbouring subtly different mutations within the same region would facilitate the further study of cis-element and microRNA clusters.

**GRAPHICAL ABSTRACT:** 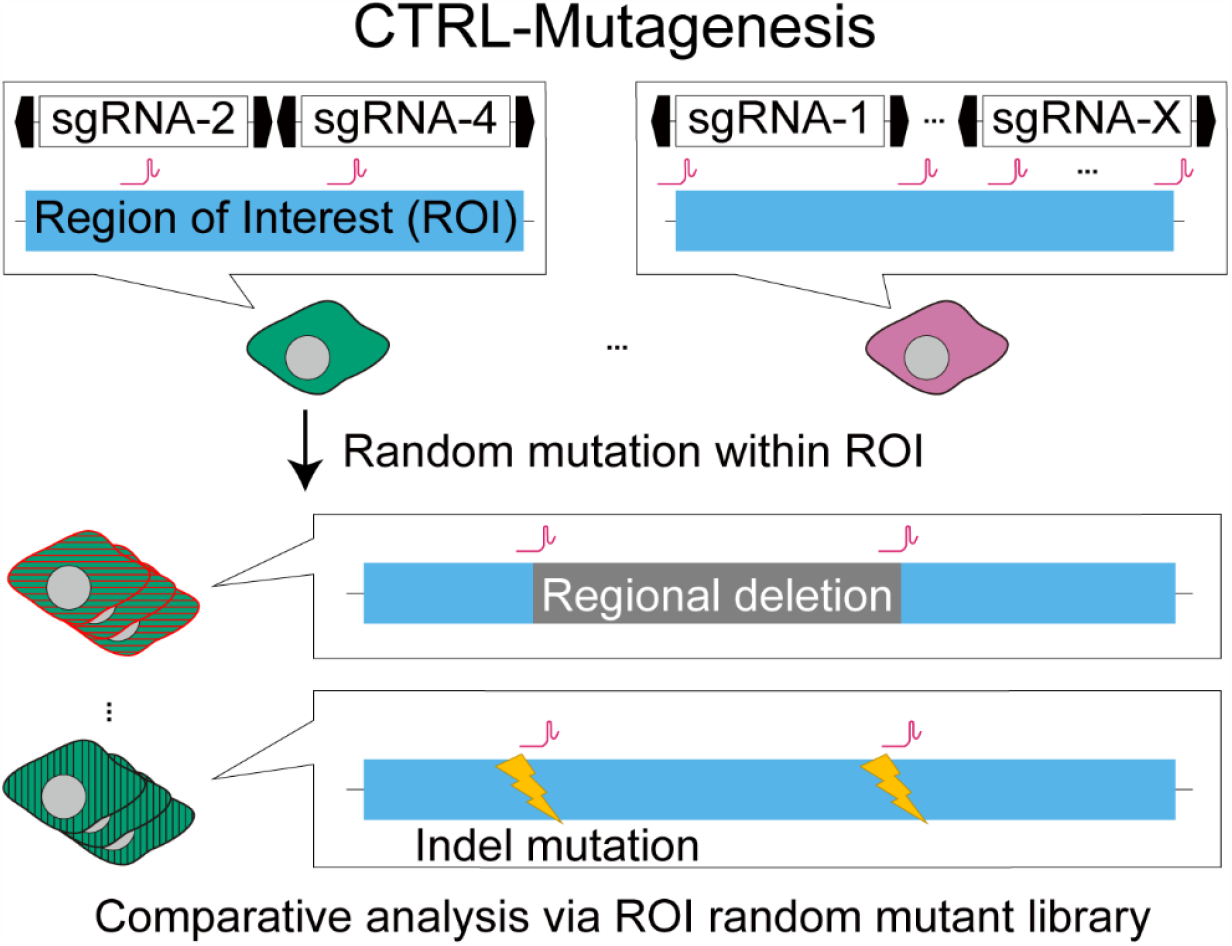

## INTRODUCTION

Whole-genome sequencing projects in humans [1], mice [2], and other species have inspired researchers to further explore how genomes regulate biological phenomena. Functional elements present within the genome include protein-coding and non-coding genes and cis-regulatory elements that regulate gene expression. The most reliable approach for identifying the functional significance of such elements in biological phenomena is to observe the phenotypes that develop when a specific element has been deactivated or deleted. While loss-of-function analysis has been conducted for numerous coding genes, very limited progress has been made on non-coding genes and cis-elements. The exons, introns, and open reading frames of coding genes have been precisely determined, making it easy to induce specific mutations for functional disruption. In mice, gene targeting techniques for the disruption of a single target gene have been available for more than 30 years, with the emergence of CRISPR-Cas9 leading to further significant progress in targeted gene disruption [3–5]. In addition, forward genetic genome-wide random mutagenesis by X-ray, *N*-ethyl-*N*-nitrosourea (ENU), and gene trapping has elucidated the functions of an extensive list of coding genes [6–8].

While it is the protein products that directly drive biological phenomena, functional elements that regulate their expression patterns are also of considerable importance. Cis-regulatory elements, such as promoters and enhancers, and non-coding transcripts, such as microRNAs (miRNAs), modulate gene expression at the pre- and post-transcriptional levels, respectively [9]. Reporter assays are usually performed to assess the regulatory capacity of such non-coding sequences [10,11]. Recently, massively parallel reporter assays (MRPAs), such as self-transcribing active regulatory region sequencing (STARR-seq), have been conducive to the identification of enhancer regions [12]. Meanwhile, reporter assays that assess regulatory capacity alone do not provide an understanding of the impact that non-coding sequences have on a given biological phenomenon. This aspect can be addressed via loss-of-function screens of non-coding sequences under biological phenomena [13]. Annotations of cis-elements have been stored in databases, such as the Encyclopedia of DNA Elements (ENCODE) [14], but these are largely predictive and not as accurate as coding gene annotations. Therefore, comparative analysis through a mutant library harbouring subtly different mutations within a target genomic region holds great promise for the identification of functionally critical regions within cis-elements. Such an approach would be effective for the functional analysis of miRNA clusters. Over 40% of human and mouse miRNA genes exist as adjacent clusters on the chromosome [15]. Several to more than a hundred miRNAs are often located in a cluster [16], and these clusters are essential for normal development and pathogenesis [17–20]. Since miRNAs frequently exhibit functional redundance [21], determining how many miRNAs and what combination of miRNAs are present in a cluster is of considerable interest. Other important aspects are how these regulate biological phenomena and what the function of each miRNA in the cluster is. While random mutagenesis within a target region could help us investigate the above, the most common whole-genome mutagenesis approaches, such as ENU and gene trapping, almost never induce deletions of genomic regions or multiple mutations within a narrow region. Therefore, the identification and functional characterisation of regulatory regions and miRNA clusters would require a novel mutagenesis approach.

In this study, we established a novel region-specific random mutagenesis approach named <u>C</u>RISPR & <u>T</u>ransposase-based <u>R</u>egiona<u>L</u> Mutagenesis (CTRL-Mutagenesis). Further, we demonstrated that CTRL-mutagenesis randomly induces diverse mutations only within the targeted regions in murine embryonic stem (mES) cells. The generated random mutant mES clone library could facilitate further functional analyses of non-coding regulatory elements within the genome.

## MATERIAL AND METHODS

### Plasmids

The PB EGFP donor vector (PB-CMV-MCS-EF1ɑ-GFP, #PB511B-1) was purchased from System Biosciences (Palo Alto, CA). The *PiggyBac* transposase (PBase) effector vector (pCX-CAG>mPBase-PGK>HSV-TK) was constructed as follows: a custom fragment containing a Kozak sequence attached to the coding sequence (CDS) of mammalian codon-optimised PBase (mPBase) (GenBanK: EF587698.1) was synthesised by Integrated DNA Technologies (Coralville, IA). The mPBase fragment was replaced from the EGFP of pCX-EGFP [22], regulated under the control of a CAG promoter. PGK>HSV-TK was PCR-amplified from pATD, a gift from Okano, H. [23] (Addgene plasmid #141012) and inserted into the pCX-CAG>mPBase, yielding pCX-CAG>mPBase-PGK>HSV-TK. The vector map of pCX-CAG>mPBase-PGK>HSV-TK is depicted in Supplementary Figure S1.

A PB single-guide RNA (sgRNA) donor vector template (pCX-PB U6>sgRNA-BbsI entry-U6>sgRNA-EGxxFP reporter-PGK>neo) was constructed as follows: the PB-5’ inverted terminal repeat (ITR)-internal domain (ID) (PB-5’ITR-ID) and PB-3’ITR-ID were PCR-amplified from PB-U6insert, a gift from Church, G.M. [24] (Addgene plasmid #104536). The U6>sgRNA-BbsI entry containing BbsI sites, which enables directional cloning of sgRNA oligos, was PCR-amplified from pX330-mC deposited in the Riken BioResource Research Center (RDB14406). The PB-5’ITR-ID, U6>sgRNA-BbsI entry, U6>sgRNA-EGxxFP reporter, PGK>neo, and PB-3’ITR-ID from the EGxxFP cassette of pCX-EGxxFP were then replaced, yielding pCX-PB U6>sgRNA-BbsI entry-U6>sgRNA-EGxxFP-PGK>neo. The vector map of pCX-PB U6>sgRNA-BbsI entry-U6>sgRNA-EGxxFP-PGK>neo is shown in Supplementary Figure S2.

PB sgRNA donor vectors targeting each *Mirc56_X*, except for *Mirc56_11*, were constructed by ligating oligos into the BbsI site of pCX-PB U6>sgRNA-BbsI entry-U6>sgRNA-EGxxFP-PGK>neo. For *Mirc56_11*, pCX-PB U6>sgRNA-*Mirc56_11*_upstream-U6>sgRNA-*Mirc56_11*_downstream-PGK>neo was constructed from the PB sgRNA donor vector.

The EGxxFP reporter vector (pCX-CAG>EGxxFP reporter-PGK>HSV-TK) was constructed as follows: the non-Chordata fragment (inverted N-part of tdTomato) containing the sgRNA-EGxxFP reporter target sequence was placed between the N and C parts of the EGFP fragment of pCX-EGxxFP [25] (Addgene plasmid #50716). PGK>HSV-TK was then inserted downstream of the CAG>EGxxFP reporter, yielding the pCX-CAG>EGxxFP reporter-PGK>HSV-TK. The vector map of pCX-CAG>EGxxFP reporter-PGK>HSV-TK is illustrated in Supplementary Figure S3.

The Cas9 editor vector (pX330-mC-PGK>HSV-TK) was constructed as follows: the U6>sgRNA scaffold cassette was excised from pX330-mC carrying human codon-optimised *Streptococcus pyogenes* Cas9 (hSpCas9)-Cdt1 (mouse) fusion protein [26]. Then, PGK>HSV-TK was inserted into the pX330-mC excised U6>sgRNA scaffold cassette, yielding pX330-mC-PGK>HSV-TK. The vector map of pX330-mC-PGK>HSV-TK is depicted in Supplementary Figure S4.

### Cells

WT mouse embryonic fibroblasts (MEFs) and neo-resistance MEFs were established from WT Crl:CD1 (ICR) or ICR^neo^ [27] foetuses at 14 d post-coitus and inactivated using 10 μg/mL mitomycin C (#M0503; Merck KGaA, Darmstadt, Germany). MEFs were cultured in the primary medium [Dulbecco’s modified Eagle’s medium (DMEM) (#11995065; Thermo Fisher Scientific, Waltham, MA) supplemented with 10% foetal bovine serum (FBS)]. C57BL/6J-derived B6J-S1^*UTR*^ mES cells [28], deposited in the Riken BioResource Cell bank (AES0140), were cultured on a feeder layer of mitomycin C-inactivated confluent WT ICR or ICR^neo^ MEFs in culture medium [DMEM supplemented with 20% KnockOut Serum Replacement (KSR) (#10828028; Thermo Fisher Scientific), 1% non-essential amino acids (#1681049; MP Biomedicals, Irvine, CA), 0.1 mM 2-mercaptoethanol (#M6250; Merck KGaA), and ESGRO-2i Supplement Kit (1:1000, #ESG1121; Merck KGaA)] in a humidified incubator at 37 °C, under 5% CO_2_ and 95% air.

Transfections were conducted as follows: for evaluation of *PiggyBac* transposon (PB transposon) integration, cells were transfected with 200 or 2000 ng of PB EGFP donor vector and 0, 350, 3500, or 17500 ng of mPBase effector vector. For the integration of the sgRNA cassette, a PB sgRNA donor vector library was prepared as a mixture of 17 PB sgRNA donor vectors (118 ng each). Cells were transfected with both 2000 ng of the PB sgRNA donor vector library and 350 ng of the mPBase effector vector. For constructing a *Mirc56* random mutant mES clone library, cells were transfected with both 2000 ng of EGxxFP reporter vector and 1000 ng of Cas9 editor vector. Cells were harvested using 0.25% trypsin-EDTA (#25200072; Thermo Fisher Scientific) and resuspended in a culture medium with 20% KSR replaced with 20% FBS. Single mES cells (7 × 10^5^ cells) were transfected with vectors using Lipofectamine LTX (#15338100; Thermo Fisher Scientific) [DNA : Lipofectamine = 1 μg : 4 μL] in 300 μL of Opti-MEM I Reduced Serum Medium (#31985062; Thermo Fisher Scientific) and then plated in 2 mL of culture medium onto a six-well plate.

Chemical selections were conducted as follows: G418 selection was conducted for 6 days in 150 μg/mL G418/culture medium (#11811023; Thermo Fisher Scientific) 2 days after transfection, as described previously [29]. Ganciclovir selection was conducted for 4 days under 10 μM ganciclovir/culture medium (#sud-gcv; InvivoGen, San Diego, CA) after G418 selection or 8 days after single-cell sorting, as described previously [30].

### Animals

An ICR outbred strain was purchased from Jackson Laboratory Japan, Inc. (Yokohama, Japan). All ICR and ICR^neo^ mice were housed in plastic cages under pathogen-free conditions in a room maintained at 23.5 °C ± 2.5 °C and 52.5% ± 12.5% relative humidity under a 14-h light:10-h dark cycle. The mice had free access to commercial chow (MF; Oriental Yeast, Tokyo, Japan) and filtered water. Animal experiments were carried out in a humane manner with approval from the Institutional Animal Experiment Committee of the University of Tsukuba, in accordance with the Regulations for Animal Experiments of the University of Tsukuba and Fundamental Guidelines for Proper Conduct of Animal Experiments and Related Activities in Academic Research Institutions under the jurisdiction of the Ministry of Education, Culture, Sports, Science, and Technology of Japan.

### Cell sorting

Cells were harvested using 0.25% trypsin-EDTA and resuspended in a culture medium with 20% KSR replaced with 20% FBS. The cell suspension was filtered through 35-μm cell strainers. The samples were then analysed on a BD FACSMelody cell sorter (BD Biosciences, Franklin Lakes, NJ), and single mES cells were sorted (Supplementary Figure S5). For single-cell cloning, the EGxxFP reporter vector and Cas9 editor vector were transfected into mES cells 2 days before fluorescence-activated cell sorting (FACS). EGFP-positive mES cells were placed onto a feeder layer of mitomycin C-inactivated WT ICR MEFs on 96-well plates. Data analysis was performed using BD FACSChorus software version 1.3.2.

### sgRNA design

To induce mutations in *Mirc56* genomic regions, we designed 16 sgRNAs (named sgRNA-*Mirc56_X)*, targeting each mature-miRNA genomic sequence, except for *Mir881* (*Mirc56_11*). Since *Mirc56_11* has no PAM sequence for hSpCas9, we designed two sgRNAs for *Mirc56_11*, with one target site upstream of *Mirc56_11* and the other downstream. The sequences of sgRNAs are listed in Supplementary Table S1.

To report both Cas9 and sgRNA expression in mES cells using the EGxxFP system [25], we designed reporter sgRNA (named sgRNA-EGxxFP reporter) targeting the non-Chordata sequence, tdTomato (Supplementary Table S1).

### Genomic DNA extraction

For the isolation of genomic DNA, mES cells were cultured on 0.1% gelatin-treated 12-well plates without MEFs. Genomic DNA was extracted via proteinase K lysis/ethanol precipitation. To this end, mES cells were lysed in lysis buffer [100 mM NaCl, 50 mM Tris-HCl (pH 8.0), 100 mM EDTA, 1% sodium dodecyl sulphate, and 6 mg/mL proteinase K]. After lysis, one-fifth 5 M NaCl was added into the lysate and then centrifuged at 13000 rpm for 5 min at 4 °C. The supernatants were added to 70% ethanol, and the genomic DNA was precipitated.

### Genotyping and short-read sequencing

Genotyping PCR was performed using AmpliTaq Gold 360 DNA Polymerase (#4398886; Thermo Fisher Scientific); the primers used are listed in Supplementary Table S2. About 200 bp was amplified around *Mirc56_X* sites. A short-read library was prepared as described previously [31], using KOD -Plus-Neo DNA Polymerase (#KMM-401; TOYOBO, Osaka, Japan) or AmpliTaq Gold 360 DNA Polymerase. The primers used for this are listed in Supplementary Table S2. Nested PCR for adding the barcode sequence was performed using relevant primers whose barcode sequences were added to the 5′ end (Supplementary Table S2). Nested PCR amplicons were purified using 1.12 × AMPure XP beads (#A63881; Beckman Coulter Genomics, Brea, CA). A 10% spike-in of PhiX control V3 (#FC-110-3001; Illumina, San Diego, CA) was added to these amplicons. Paired-end sequencing (2 × 150 bases) of these amplicons was performed using iSeq 100 i1 Reagent v2 (#20031371; Illumina) and iSeq 100 (Illumina).

Sequencing reads were de-multiplexed using the GenerateFASTQ module version 2.0.0 on iSeq 100 Software (Illumina). Analysis of on-target amplicon sequencing was performed using CRISPResso2 version 2.2.9 in batch mode [32].

## RESULTS

### Scheme of CTRL-Mutagenesis

We developed a novel CTRL-Mutagenesis method for inducing random and diverse mutations only within a region of interest (ROI). The overall scheme of CTRL-Mutagenesis is illustrated in Figure 1. A DNA vector library expressing each sgRNA for the respective target sites in the ROI is introduced into cultured cells via *PiggyBac* transposase (PBase). To evaluate the *in vivo* and *in vitro* feasibility of this method, we selected mES cells. We obtained mES cells with various combinations of sgRNA expression cassettes integrated, naming these PB mES cells. To induce random mutations within the ROI, PB mES cells are treated with Cas9, which cleaves target sequences guided by the randomly integrated sgRNA. Multiple cleavages of cis-sequences induce insertion and/or deletion (Indel) mutations at each cleaved site and/or regional deletions flanked by cleaved sites [33]. Therefore, even if the PB mES cells carry an identical combination of sgRNA cassettes, diverse mutations are induced within the ROI. A random combination of sgRNA cassettes integrated via the *PiggyBac* system and random on-target CRISPR-Cas9 mutagenesis enhance the randomness of mutation combinations induced. As a result, CTRL-Mutagenesis yields an ROI random mutant mES clone library after single-cell cloning.

**Figure 1.**
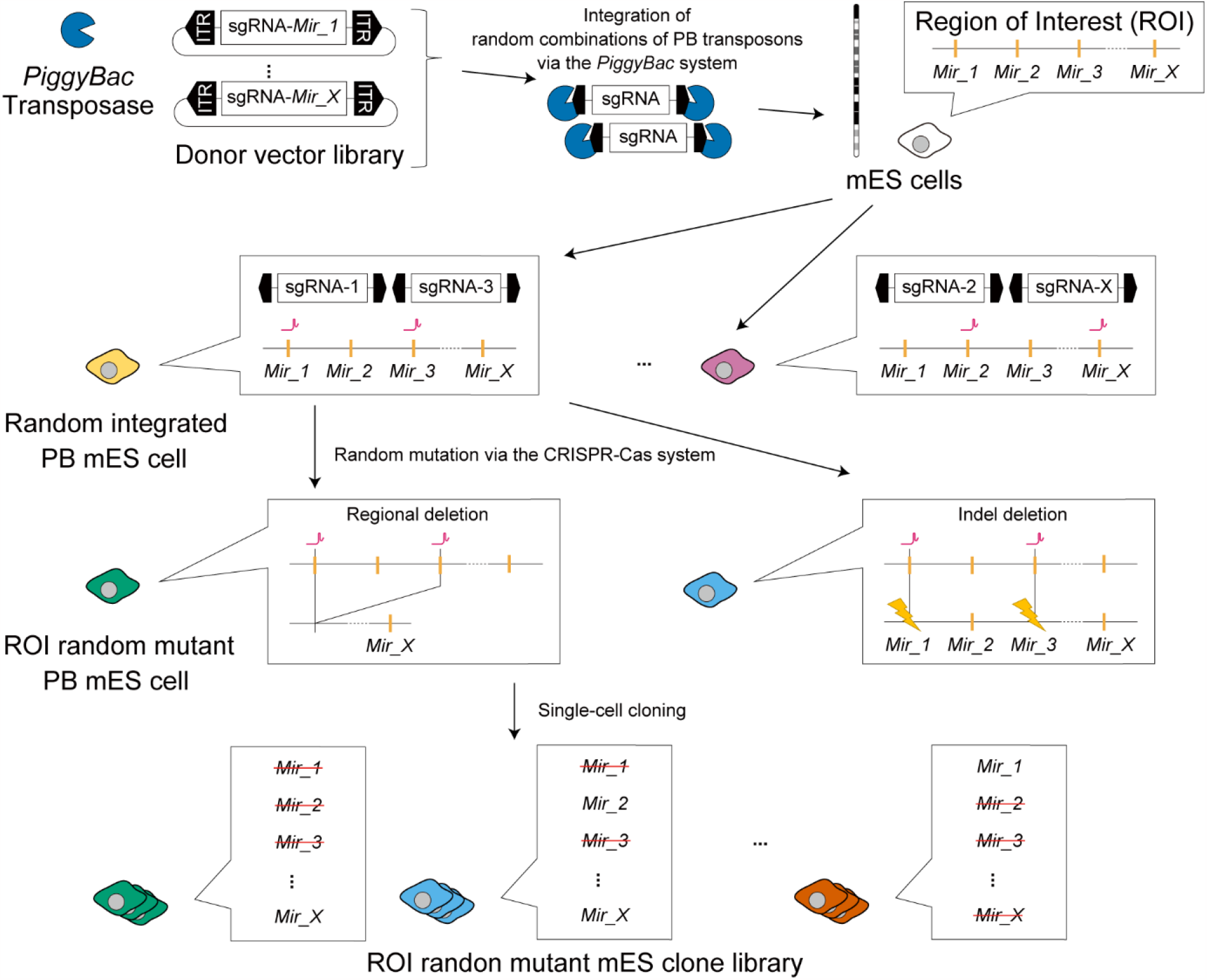
Scheme of <u>C</u>RISPR & <u>T</u>ransposase based <u>R</u>egiona<u>L</u> Mutagenesis (CTRL-Mutagenesis). Magenta symbols indicate sgRNA target sites. Yellow lightning symbols indicate Indel mutations.

### Evaluation of *PiggyBac* transposon integration

CTRL mutagenesis requires the efficient introduction of multiple sgRNA expression cassettes into the chromosomes of host cells. To determine optimal doses, we transfected cells with 200 or 2000 ng of donor vector carrying the *EGFP* expression cassette as well as with 0, 350, 3500, or 17500 ng of effector vector carrying mammalian codon-optimised PBase (mPBase) [34,35] expression cassettes into mES cells (Figure 2A). Without effector vector transfection, transient expression achieved with 200 ng of the donor vector lasted 6 days after transfection, while that from 2000 ng of the donor vector lasted for 8 days (Figure 2B). Thus, we defined EGFP expression after 10 days as an indicator of stable gene transduction. At both donor vector concentrations, mES cells transfected with 350 ng of effector vectors showed the highest EGFP-positive ratio 10 days after transfection. Besides, a very high concentration (17500 ng) of effector vectors induced the lowest EGFP-positive ratio among mES cells transfected with effector vectors. We then evaluated the intensity of EGFP signals (Figure 2C). The mES cells transfected with 2000 ng of donor and 350 ng of effector vectors showed broader EGFP intensity than did those transfected with 200 ng of donor and 350 ng of effector vectors. EGFP signal intensity correlated with the copy number of EGFP cassettes integrated into genomes [36]. These results suggested that even transfection with a higher dose of donor vector could integrate low to high copy numbers of donor vectors into the genome. Thus, we decided to use 2000 ng of donor and 350 ng of effector vectors for future analyses since CTRL-mutagenesis requires diverse sgRNA expression vectors.

**Figure 2.**
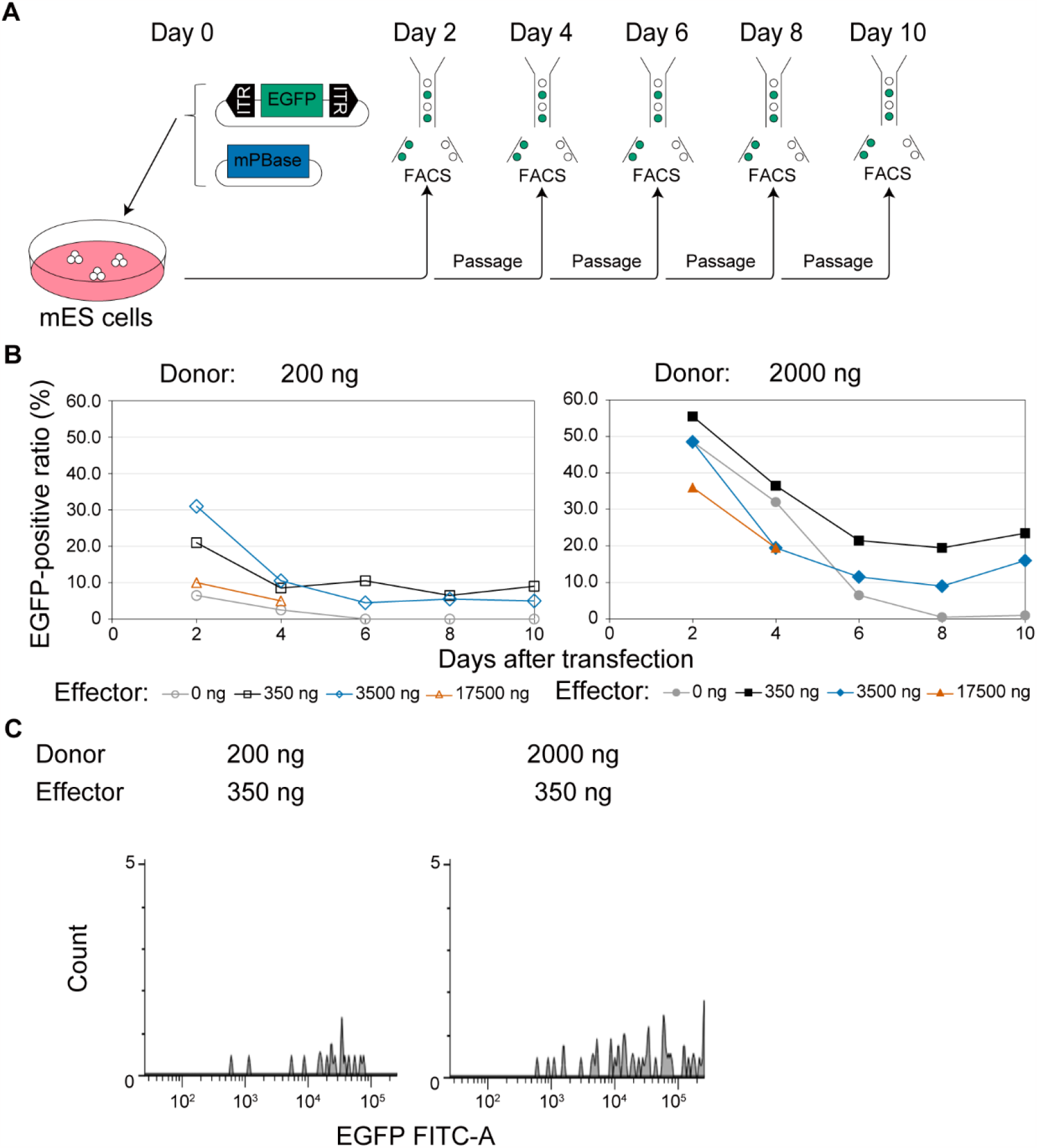
Evaluation of *PiggyBac* transposon integration. **A**, Time course for evaluation of *PiggyBac* transposon (PB transposon) integration. The day when the donor vector carrying EGFP flanked by *PiggyBac* inverted terminal repeats (ITRs) and the effector vector carrying mammalian codon-optimised *PiggyBac* transposase (mPBase) were transfected at several concentrations was defined as Day 0. Fluorescence-activated cell sorting (FACS) was conducted every 2 days after transfection to confirm the ration of EGFP-positive mES cells. Passages were conducted using the remaining murine embryonic stem (mES) cells after FACS. **B**, Ratio of EGFP-positive mES cells and days after transfection. EGFP-positive mES cells were calculated among a total of 200 mES cells passing quality filters by FACS. Blank symbols in the left figure indicate mES cells transfected with 200 ng of donor vector, and filled symbols in the right figure indicate those transfected with 2,000 ng of donor vector. Gray circles, black squares, blue diamonds, and orange triangles indicate mES cells transfected with 0, 350, 3500, and 17500 ng of effector vector, respectively. We stopped evaluating mES cells transfected with a very high concentration (17500 ng) of effector vector on Day 4. **C**, Histogram of EGFP-positive mES cells on Day 10. The upper table shows the conditions of transfection. Lower histograms show all EGFP-positive mES cells in a total of 200 mES cells passing quality filters. The x-axis shows EGFP signal intensity, and the y-axis shows counts of EGFP-positive mES cells.

### Construction of *Mirc56* random mutant mES clone library

To prove that CTRL-Mutagenesis randomly induces diverse mutations within the targeted ROI, we focused on miRNA cluster *Mirc56* on the X chromosome (Figure 3). There are 19 miRNAs in *Mirc56* (hereafter, each miRNA is referred to as *Mirc56_Xs*), interspersed within the 64 kb genomic region. We targeted *Mirc56* as a proof of concept for the following three reasons. First, *Mirc56* is located on the X chromosome, and we used the male B6J-S1^*UTR*^ mES cell line [28] in this study, which allows for the monoallelic assessment of the genotype. Second, it is the largest miRNA cluster on the X chromosome. Third, *Mirc56* is not expressed in mES cells, and mutating it probably does not affect survival or proliferation. To induce mutations in *Mirc56_X*s, we constructed a sgRNA donor vector library carrying a neo-resistance gene and two sgRNAs, one targeting each *Mirc56_X* (Figure 3) and the other targeting EGxxFP. We transfected 2000 ng of the sgRNA donor vector library and 350 ng of the mPBase effector vector carrying mPBase and HSV-TK into mES cells (Figure 4A). We obtained bulk PB mES cells with a chromosomally integrated sgRNA donor vector and without the mPBase effector vector via positive selection with G418 and negative selection with ganciclovir (Figure 4A). To confirm the integration of the various sgRNA donors, we performed targeted short next-generation sequencing (NGS) using bulk genomic DNA from PB mES cells as a template to sequence the sgRNAs introduced into chromosomes (Figure 4B). Targeted short NGS revealed the integration of all kinds of sgRNAs, but two (sgRNA for *Mirc56_2* and *4*) were rarely detected. To induce random mutations on *Mirc56*, we transfected the EGxxFP reporter vector and Cas9 editor vector (Figure 4A). Both vectors carried HSV-TK. To efficiently obtain *Mirc56* mutant mES cells, we performed positive selection with the EGxxFP system reporting CRISPR-Cas system activity; that is, Cas-induced cleavage of EGxxFP led to EGFP expression [25]. The negative selection was carried out using ganciclovir. The sgRNA donor vectors had sgRNAs targeting not only each *Mirc56_X* but also EGxxFP. Co-transfection of PB mES cells with a Cas9 editing vector and an EGxxFP reporter vector resulted in mutations within the *Mirc56* genomic region and the conversion of EGxxFP to EGFP. To obtain only *Mirc56* random mutant PB mES cells with an EGFP signal, we sorted single EGFP-positive PB mES cells via FACS, with the gates set to exclude PB mES cells transfected only with the EGxxFP reporter vector (Figure 4C). The ratio of EGFP-positive PB mES cells was 4.0% with Cas9 editor and EGxxFP reporter vectors. We then added ganciclovir to the medium during single-cell cloning to eliminate PB mES cells in which the editor or reporter was chromosomally integrated. Through PCR, we confirmed that 87 out of 89 clones carried no integrations of effector, editor, and reporter vectors (data not shown). Finally, we obtained a *Mirc56* random mutant PB mES clone library that consisted of 87 mutant clones.

**Figure 3.**
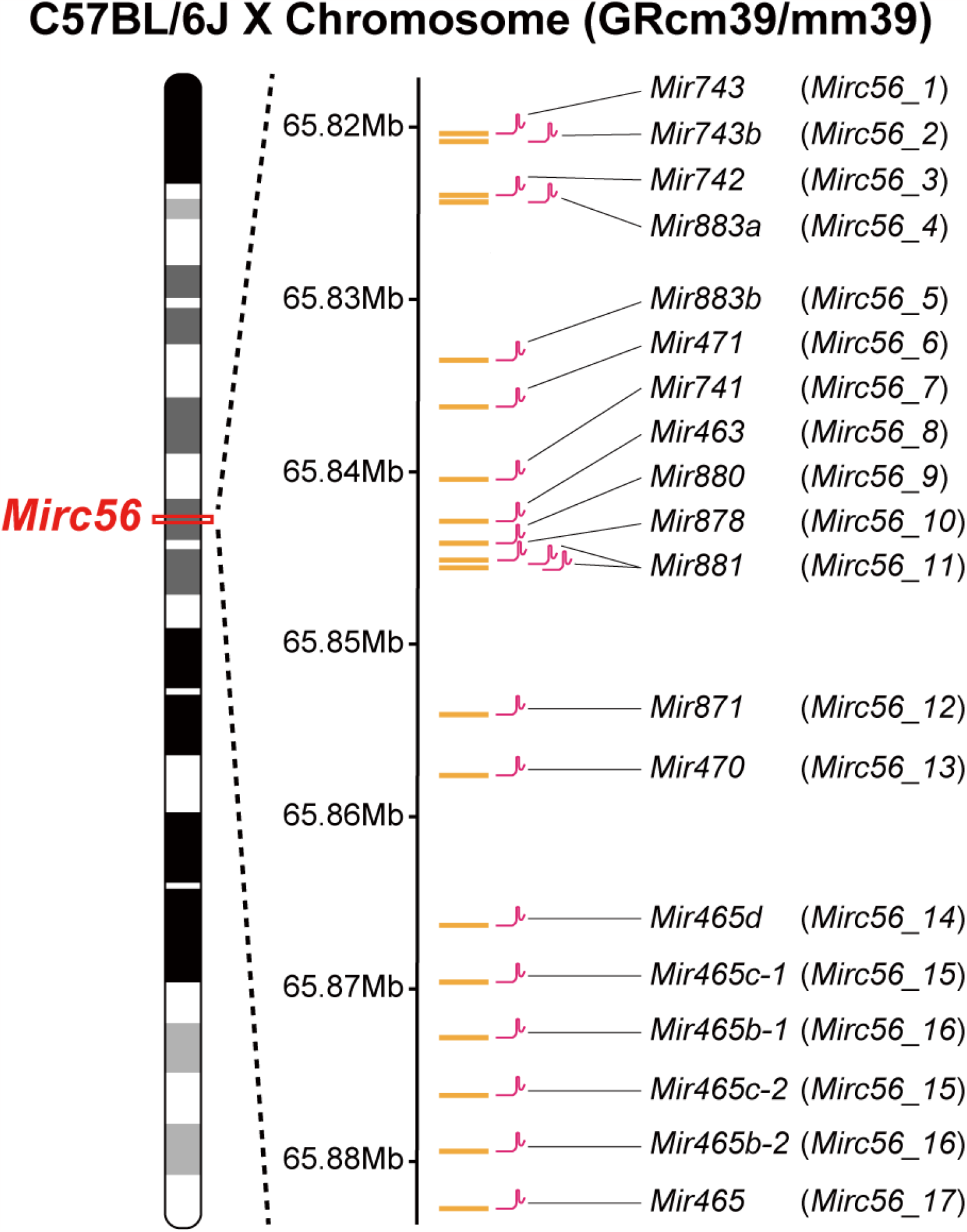
Gene map of the *Mirc56* region on the X chromosome (65,820,363-65,882,731 bp, GRCm39/mm39). *Mirc56* harbours 19 miRNAs. *Mir465c-1* and *Mir465c-2* (*Mirc56_15*), as well as *Mir465b-1* and *Mir465b-2* (*Mirc56_16*), have identical sequences at the mature-miRNA genomic regions. Magenta symbols indicate sgRNA target sites on *Mirc56_X*’s mature-miRNA genomic sequences, except for *Mir881* (*Mirc56_11*). For deletion of *Mirc56_11*, one of the two sgRNA-*Mirc56_11* targets a region upstream of *Mirc56_11*, while the other targets a downstream region.

**Figure 4.**
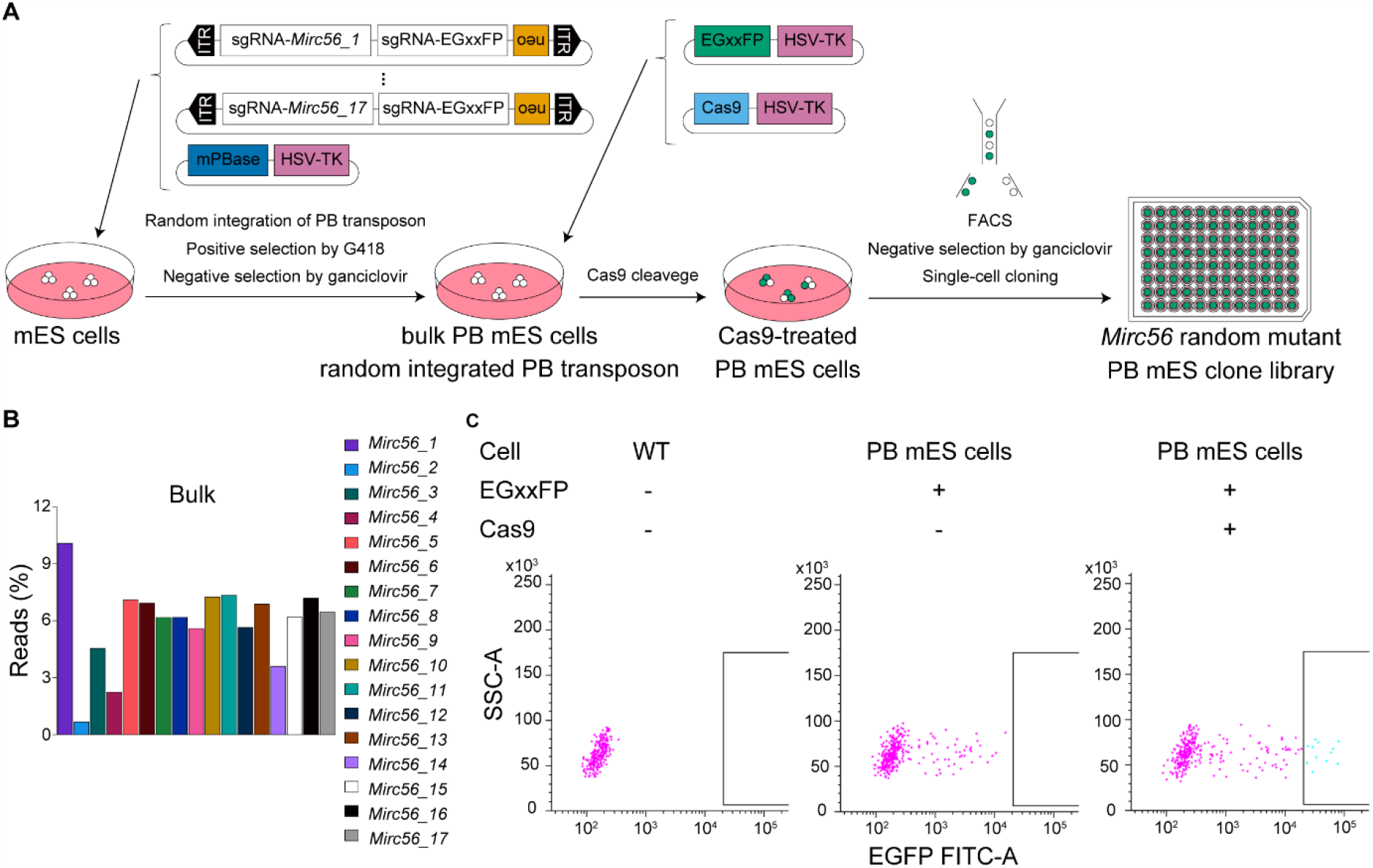
Construction of the *Mirc56* random mutant mES clone library. **A**, Workflow for library construction. **B**, Random integrated sgRNA-*Mirc56_X* cassettes in bulk PB mES cells. The bar colour shows each sgRNA-*Mirc56_X* cassette. The y-axis represents the percentage of reads with targeted amplicon short next-generation sequencing (NGS). **C**, EGFP-based single-cell sorting in bulk PB mES cells. The upper table shows the conditions of transfection. Lower scatter graphs show each mES cell passing quality filters in a total of 2000 events. The y-axis shows the EGFP signal intensity. The boxes in scatter graphs show the gates of the EGFP filter.

### Evaluation of random integration and random mutation in *Mirc56* random mutant mES clone library

To determine whether PB mES clones carried various combinations of sgRNA cassettes, we amplified sgRNA cassettes for *Mirc56_X* and conducted targeted short NGS on 87 clones (Figure 5A). This heatmap indicated that diverse combinations of sgRNA were integrated, except for the sgRNA cassettes for *Mirc56_2* and *4*. This rare integration of sgRNA cassettes for *Mirc56_2* and *4* was consistent with the trend in bulk PB mES cells (Figure 4B). Here, we focused on the number of sgRNA cassette varieties. The maximum number of sgRNA cassette varieties integrated into PB mES clones was 16, with an average of 4.7 (Figure 5B). Besides, to evaluate the properties of our *Mirc56* random mutant PB mES clone library, we determined the genotypes of *Mirc56* genomic regions, except for *Mirc56_14, 15, 16*, and *17*, in 87 clones. We skipped genotyping *Mirc56_14, 15, 16*, and *17* (*Mir465* cluster) because PCR-based genotyping within this region was difficult owing to six tandem repeats [37]. The generated mutation map indicated that almost all clones seemed to harbour different combinations of mutations (Figure 5C). There seemed to be expected mutation events in each clone; that is, a single sgRNA induced Indel mutation in its own target *Mirc56_X*, and multiple sgRNAs induced regional deletion flanked by target *Mirc56_X*s. These results suggested that Indel mutations and regional deletions were sgRNA-dependent. Besides, the complex combinations of regional deletions and Indel mutations suggested that Cas9 could induce multiple mutation events on the same strand. To further evaluate random mutations by Cas9, we focused on *Mirc56_Xs* targeted by the expressed sgRNA in each clone (Figure 5D). We excluded *Mirc56_11* from this evaluation because it was targeted by two sgRNAs (Figure 3). Regional deletions were dominant. In addition, an average of 22.7 *Mirc56_X* sites were targeted in 87 clones, and the frequency of target sites was similar with each *Mirc56_X*, except for *Mirc56_2* and *4*. These results indicated that almost all kinds of sgRNA cassettes could integrate with the same frequency except for the *Mirc56_2*- and *4*-targeting cassettes. Next, we investigated the mutation rate at each target *Mirc56_X* site. On average, 80.5% of target sites had Indel mutations or regional deletions. These results suggested that CTRL-Mutagenesis could induce diverse mutations within target sites.

**Figure 5.**
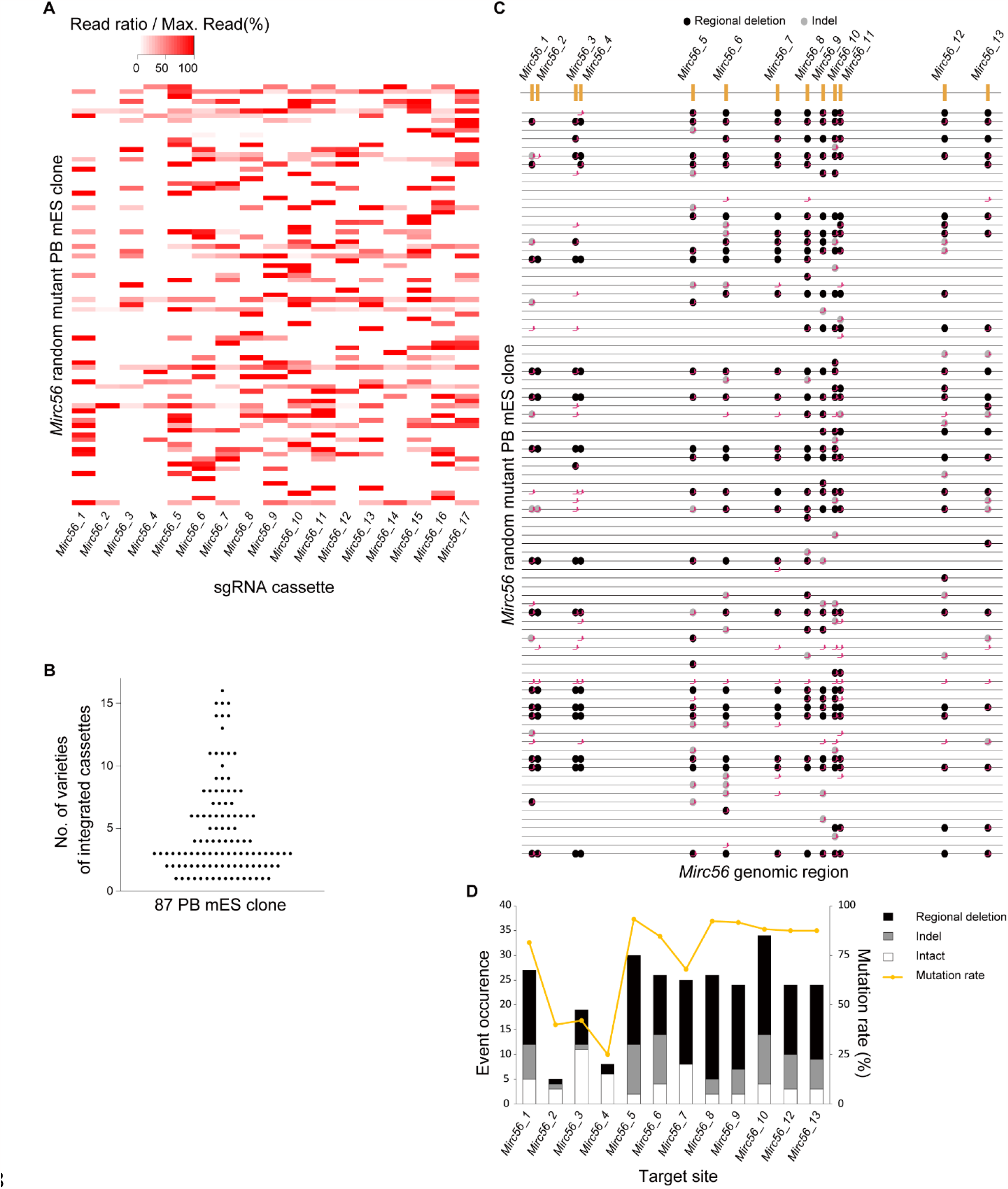
Evaluation of random integration and random mutation in the *Mirc56* random mutant mES clone library. **A**, Heatmap of integrated sgRNA cassettes in 87 *Mirc56* random mutant clones. The read counts of each sgRNA cassette were divided by the maximum read counts in each *Mirc56* random mutant clone. **B**, Scatter graph with the number of varieties of integrated cassettes in 87 *Mirc56* random mutant clones. **C**, Genomic map of *Mirc56* in 87 *Mirc56* random mutant clones. The x-axis shows the genomic region of *Mirc56_1, 2, 3, 4, 5, 6, 7, 8, 9, 10, 11, 12*, and *13* without *Mirc56_14, 15, 16*, and *17*. Magenta symbols indicate sgRNA target sites in each PB mES clone. **D**, Mutations in target sites in 87 *Mirc56* random mutant clones. The target sites do not include *Mirc56_11, 14, 15, 16*, and *17*. The left vertical axis and bar graphs show event occurrence, while the right vertical axis and line graph show the mutation rate on target sites in 87 *Mirc56* random mutant clones. The bar colour indicates each event (Black: Regional deletion, Gray: Indel mutation, White: Intact).

## DISCUSSION

Comparative analysis of a mutant library harbouring subtly different mutations within the same region is particularly useful for the functional analysis of non-coding sequences. In this report, we introduce an approach for ROI-targeted random mutagenesis. Further, we demonstrated that our novel method, named CTRL-Mutagenesis, could randomly induce diverse mutations within the target ROI. A random combination of sgRNA cassettes integrated using the *PiggyBac* system and random on-target CRISPR-Cas9 mutagenesis was employed to enhance the randomness of induced mutation combinations.

CTRL-Mutagenesis was employed for efficiently constructing an ROI random mutant library. To this end, we utilised three selection systems, namely, G418 selection [29], ganciclovir selection [30], and the EGxxFP system [25]. These allowed for the specific selection of ROI random mutant PB mES clones without the integration of unintended vectors. In fact, 87 out of 89 evaluated clones carried no unintended vectors. Targeted short NGS captured the integration of various sgRNA cassettes in these 87 clones. We expected that targeted short NGS could identify precise combinations of integrated sgRNA cassettes because Indel mutations and regional deletions depended on integrated sgRNA cassettes. As a result, an ROI random mutant library consisting of 87 clones was efficiently constructed and evaluated.

To optimise the integration of sgRNA cassettes, we should improve two conditions. First is the integration frequency among sgRNA cassettes. Our conditions allowed for the integration of sgRNA cassettes at the same frequency, except for sgRNA cassettes targeting *Mirc56_2* and *4* (Figures 5A and 5D). The lower frequencies noted in the case of these cassettes were also observed in bulk PB mES cells (Figure 4B). We considered that such lower frequencies were derived from the imbalanced integration into bulk PB mES cells rather than from the imbalanced selection of PB mES clones via FACS. We suspect this was caused by a technical error, such as an unequal amount of sgRNA donor vector or the sequence in sgRNA cassettes affecting integration efficiency or cell growth. The second condition is the number of integrated sgRNA cassettes. Out of 17 kinds of donor vectors, our conditions integrated 4.7 on average (Figure 5B). The dose of the donor vector had a greater impact on PB transposon integration into genomes than did the effector vector dose (Figure 2B), in accordance with a previous report [29]. In addition, we revealed that even transfection with a higher dose of donor vector resulted in the integration of low to high copy numbers into genomes (Figure 2C). Therefore, higher doses of donor vector should further improve integration copy number and efficiency. Tuning the dose of the donor vector or employing another PBase, such as hyperactive PBase (hyPBase) [35], could control the number of sgRNA cassettes to be integrated. Of note, off-target effects could comparatively affect phenotypes because of high risk that is further increased by two factors. The first factor is the integration of the PB transposon into random TTAA sites across genomes [38]. To avoid this, one of solutions is to use excision-only-PBase [39] to remove the PB transposon from the ROI random mutant library. The second factor is mutations in the non-specific mismatch genomic sites of sgRNAs. The risk increases along with the number of sgRNA varieties. Alternatively, conventional methods using minimum sgRNAs to regenerate functionally critical region mutants showing phenotypes reduce the risk.

CTRL-Mutagenesis could randomly induce diverse mutations. However, regional deletion was dominant in our mutant library (Figure 5C, D). One explanation is that *Mirc56_X* sites were deleted by flanked cleavage sites even if either they were not the target sites or Indel mutations were induced at target sites. This is one of our limitations in constructing a mutant library that harbours subtly different mutations within the same region. One way to overcome this is using a mixture of Cas9 and base editor [40] or Cas nickase and gRNA designed on the same strand [41]. These options should induce Indel mutation or substitutions while avoiding or reducing regional deletion.

In the present study, we subjected mES cells to CTRL-Mutagenesis. To validate the efficacy of our method, further comparative analysis under *in vitro* and *in vivo* conditions remains to be performed. We selected mES cells as these can be used to recapitulate biological development *in vitro* and *in vivo* [28,42]. Organoids are one example of a rapid evaluation system for such functional analysis [43]. In this study, we targeted a miRNA cluster genomic region, but we also propose the application of CTRL-Mutagenesis for targeting other non-coding chromosomal regions such as enhancers or promoters. This would require CTRL-Mutagenesis to induce regional deletions at the respective regions. Taken together, our CTRL-Mutagenesis is expected to be of great value *in vitro* and *in vivo* comparative analyses with the aim of elucidating the functional importance of non-coding regions.

## DATA AVAILABILITY

All sequence data (FASTQ) (Illumina iSeq) from this study can be retrieved from Sequence Read Archive (SRA) under the BioProject ID PRJNA996747. The custom R and Bash scripts used to quantify the amounts of 17 sgRNA cassettes for each mES clone are available at DOI: 10.6084/m9.figshare.24046311. The custom R script to produce the heatmap of sgRNA cassette integration coverages can be downloaded from DOI: 10.6084/m9.figshare.24046314.

## SUPPLEMENTARY DATA

Supplementary Data are available at bioRxiv.

## ACKNOWLEDGEMENTS

We would like to thank Editage (www.editage.com) for English language editing.

## REFERENCES

1. Abdellah Z, Ahmadi A, Ahmed S, Aimable M, Ainscough R, Almeida J, et al. Finishing the euchromatic sequence of the human genome. Nature. 2004;431: 931–945. doi:10.1038/nature03001

2. Waterston RH, Lindblad-Toh K, Birney E, Rogers J, Abril JF, Agarwal P, et al. Initial sequencing and comparative analysis of the mouse genome. Nature. 2002;420: 520–562. doi:10.1038/nature01262

3. Kohan DE. Progress in gene targeting: using mutant mice to study renal function and disease. Kidney Int. 2008;74: 427–437. doi:10.1038/KI.2008.146

4. Capecchi MR. Gene targeting in mice: Functional analysis of the mammalian genome for the twenty-first century. Nat Rev Genet. 2005;6: 507–512. doi:10.1038/nrg1619

5. Zhang HX, Zhang Y, Yin H. Genome Editing with mRNA Encoding ZFN, TALEN, and Cas9. Mol Ther. 2019;27: 735–746. doi:10.1016/J.YMTHE.2019.01.014

6. Thomas JW, Lamantia C, Magnuson T. X-ray-induced mutations in mouse embryonic stem cells. Proc Natl Acad Sci U S A. 1998;95: 1114–1119. doi:10.1073/PNAS.95.3.1114/ASSET/52C4E597-9D49-4977-8489-61A106585FD7/ASSETS/GRAPHIC/PQ0383986004.JPEG

7. Justice MJ, Noveroske JK, Weber JS, Zheng B, Bradley A. Mouse ENU Mutagenesis. Hum Mol Genet. 1999;8: 1955–1963. doi:10.1093/HMG/8.10.1955

8. Stanford WL, Cohn JB, Cordes SP. Gene-trap mutagenesis: Past, present and beyond. Nat Rev Genet. 2001;2: 756–768. doi:10.1038/35093548

9. Pagni S, Mills JD, Frankish A, Mudge JM, Sisodiya SM. Non-coding regulatory elements: Potential roles in disease and the case of epilepsy. Neuropathol Appl Neurobiol. 2022;48: e12775. doi:10.1111/NAN.12775

10. Dong ZQ, Hu ZG, Li HQ, Jiang YM, Cao MY, Chen P, et al. Construction and characterization of a synthetic Baculovirus-inducible 39K promoter. J Biol Eng. 2018;12: 1–12. doi:10.1186/S13036-018-0121-8/FIGURES/5

11. Sharma CM, Vogel J. Experimental approaches for the discovery and characterization of regulatory small RNA. Curr Opin Microbiol. 2009;12: 536–546. doi:10.1016/J.MIB.2009.07.006

12. Muerdter F, Boryń ŁM, Arnold CD. STARR-seq — Principles and applications. Genomics. 2015;106: 145–150. doi:10.1016/J.YGENO.2015.06.001

13. Ohmura S, Mizuno S, Oishi H, Ku CJ, Hermann M, Hosoya T, et al. Lineage-affiliated transcription factors bind the Gata3 Tce1 enhancer to mediate lineage-specific programs. J Clin Invest. 2016;126: 865–878. doi:10.1172/JCI83894

14. Dunham I, Kundaje A, Aldred SF, Collins PJ, Davis CA, Doyle F, et al. An integrated encyclopedia of DNA elements in the human genome. Nature. 2012;489. doi:10.1038/nature11247

15. Griffiths-Jones S, Saini HK, Van Dongen S, Enright AJ. miRBase: Tools for microRNA genomics. Nucleic Acids Res. 2008;36: 154–158. doi:10.1093/nar/gkm952

16. Sun J, Gao B, Zhou M, Wang Z zhen, Zhang F, Deng J en, et al. Comparative genomic analysis reveals evolutionary characteristics and patterns of microRNA clusters in vertebrates. Gene. 2013;512: 383–391. doi:10.1016/J.GENE.2012.09.102

17. Ventura A, Young AG, Winslow MM, Lintault L, Meissner A, Erkeland SJ, et al. Targeted Deletion Reveals Essential and Overlapping Functions of the miR-17∼92 Family of miRNA Clusters. Cell. 2008;132: 875–886. doi:10.1016/j.cell.2008.02.019

18. Hébert SS, Horré K, Nicolaï L, Papadopoulou AS, Mandemakers W, Silahtaroglu AN, et al. Loss of microRNA cluster miR-29a/b-1 in sporadic Alzheimer’s disease correlates with increased BACE1/β-secretase expression. Proc Natl Acad Sci U S A. 2008;105: 6415–6420. doi:10.1073/PNAS.0710263105/SUPPL_FILE/SD3.XLS

19. Besser J, Malan D, Wystub K, Bachmann A, Wietelmann A, Sasse P, et al. MiRNA-1/133a Clusters Regulate Adrenergic Control of Cardiac Repolarization. PLoS One. 2014;9: e113449. doi:10.1371/JOURNAL.PONE.0113449

20. Zhou L, Miller C, Miraglia LJ, Romero A, Mure LS, Panda S, et al. A genome-wide microRNA screen identifies the microRNA-183/96/182 cluster as a modulator of circadian rhythms. Proc Natl Acad Sci U S A. 2021;118: e2020454118. doi:10.1073/PNAS.2020454118/SUPPL_FILE/PNAS.2020454118.SAPP.PDF

21. Fischer S, Handrick R, Aschrafi A, Otte K. Unveiling the principle of microRNA-mediated redundancy in cellular pathway regulation. RNA Biol. 2015;12: 238–247. doi:10.1080/15476286.2015.1017238/SUPPL_FILE/KRNB_A_1017238_SM0586.ZIP

22. Okabe M, Ikawa M, Kominami K, Nakanishi T, Nishimune Y. “Green mice” as a source of ubiquitous green cells. FEBS Lett. 1997;407: 313–319. doi:10.1016/S0014-5793(97)00313-X

23. Yoshimatsu S, Sone T, Nakajima M, Sato T, Okochi R, Ishikawa M, et al. A versatile toolbox for knock-in gene targeting based on the Multisite Gateway technology. PLoS One. 2019;14: e0221164. doi:10.1371/JOURNAL.PONE.0221164

24. Kalhor R, Kalhor K, Mejia L, Leeper K, Graveline A, Mali P, et al. Developmental barcoding of whole mouse via homing CRISPR. Science. 2018;361. doi:10.1126/science.aat9804

25. Mashiko D, Fujihara Y, Satouh Y, Miyata H, Isotani A, Ikawa M. Generation of mutant mice by pronuclear injection of circular plasmid expressing Cas9 and single guided RNA. Sci Rep. 2013;3: 1–6. doi:10.1038/srep03355

26. Mizuno-Iijima S, Ayabe S, Kato K, Matoba S, Ikeda Y, Dinh TTH, et al. Efficient production of large deletion and gene fragment knock-in mice mediated by genome editing with Cas9-mouse Cdt1 in mouse zygotes. Methods. 2021;191: 23–31. doi:10.1016/j.ymeth.2020.04.007

27. Takahashi S, Onodera K, Motohashi H, Suwabe N, Hayashi N, Yanai N, et al. Arrest in Primitive Erythroid Cell Development Caused by Promoter-specific Disruption of the GATA-1 Gene. J Biol Chem. 1997;272: 12611–12615. doi:10.1074/JBC.272.19.12611

28. Tanimoto Y, Iijima S, Hasegawa Y, Suzuki Y, Daitoku Y, Mizuno S, et al. Embryonic Stem Cells Derived from C57BL / 6J and C57BL / 6N Mice. Comp Med. 2008;58: 347–352.

29. Wang W, Lin C, Lu D, Ning Z, Cox T, Melvin D, et al. Chromosomal transposition of PiggyBac in mouse embryonic stem cells. Proc Natl Acad Sci U S A. 2008;105: 9290–9295. doi:10.1073/pnas.0801017105

30. Naujok O, Kaldrack J, Taivankhuu T, Jörns A, Lenzen S. Selective Removal of Undifferentiated Embryonic Stem Cells from Differentiation Cultures Through HSV1 Thymidine Kinase and Ganciclovir Treatment. Stem Cell Rev Reports. 2010;6: 450–461. doi:10.1007/s12015-010-9148-z

31. Tamari T, Ikeda Y, Morimoto K, Kobayashi K, Mizuno-Iijima S, Ayabe S, et al. A universal method for generating knockout mice in multiple genetic backgrounds using zygote electroporation. Biol Open. 2023;12. doi:10.1242/BIO.059970

32. Clement K, Rees H, Canver MC, Gehrke JM, Farouni R, Hsu JY, et al. CRISPResso2 provides accurate and rapid genome editing sequence analysis. Nature Biotechnology. Nature Publishing Group; 2019. pp. 224–226. doi:10.1038/s41587-019-0032-3

33. Kraft K, Geuer S, Will AJ, Chan WL, Paliou C, Borschiwer M, et al. Deletions, inversions, duplications: Engineering of structural variants using CRISPR/Cas in mice. Cell Rep. 2015;10: 833–839. doi:10.1016/j.celrep.2015.01.016

34. Cadiñanos J, Bradley A. Generation of an inducible and optimized piggyBac transposon system. Nucleic Acids Res. 2007;35. doi:10.1093/nar/gkm446

35. Yusa K, Zhou L, Li MA, Bradley A, Craig NL. A hyperactive piggyBac transposase for mammalian applications. Proc Natl Acad Sci U S A. 2011;108: 1531–1536. doi:10.1073/pnas.1008322108

36. Kolacsek O, Krízsik V, Schamberger A, Erdei Z, Apáti Á, Várady G, et al. Reliable transgene-independent method for determining Sleeping Beauty transposon copy numbers. Mob DNA. 2011;2: 1–8. doi:10.1186/1759-8753-2-5/TABLES/2

37. Morimoto K, Numata K, Daitoku Y, Hamada Y, Kobayashi K, Kato K, et al. Reverse genetics reveals single gene of every candidate on Hybrid sterility, X Chromosome QTL 2 (Hstx2) are dispensable for spermatogenesis. Sci Rep. 2020;10: 1–9. doi:10.1038/s41598-020-65986-y

38. Yusa K. piggyBac transposon. Mob DNA III. 2015; 873–890. doi:10.1128/9781555819217.ch39

39. Wang G, Yang L, Grishin D, Rios X, Ye LY, Hu Y, et al. Efficient, footprint-free human iPSC genome editing by consolidation of Cas9/CRISPR and piggyBac technologies. Nat Protoc. 2017;12: 88–103. doi:10.1038/nprot.2016.152

40. Gaudelli NM, Komor AC, Rees HA, Packer MS, Badran AH, Bryson DI, et al. Programmable base editing of T to G C in genomic DNA without DNA cleavage. Nature. 2017;551: 464–471. doi:10.1038/nature24644

41. Gopalappa R, Suresh B, Ramakrishna S, Kim HH. Paired D10A Cas9 nickases are sometimes more efficient than individual nucleases for gene disruption. Nucleic Acids Res. 2018;46: e71–e71. doi:10.1093/nar/gky222

42. Omori H, Otsu M, Nogami H, Shibata M. Heat shock response enhanced by cell culture treatment in mouse embryonic stem cell-derived proliferating neural stem cells. PLoS One. 2021;16: e0249954. doi:10.1371/JOURNAL.PONE.0249954

43. Rossi G, Manfrin A, Lutolf MP. Progress and potential in organoid research. Nature Reviews Genetics. Nature Publishing Group; 2018. pp. 671–687. doi:10.1038/s41576-018-0051-9

