## Supplemental Figure S1 for "Regional random mutagenesis driven by multiple sgRNAs and diverse on-target genome editing events to identify functionally important elements in non-coding regions"

**MANUSCRIPT TITLE**

### TABLE LEGENDS

Supplementary Table S1. Sequence of sgRNAs used in this study.

Supplementary Table S2. Sequence of primers used in this study.

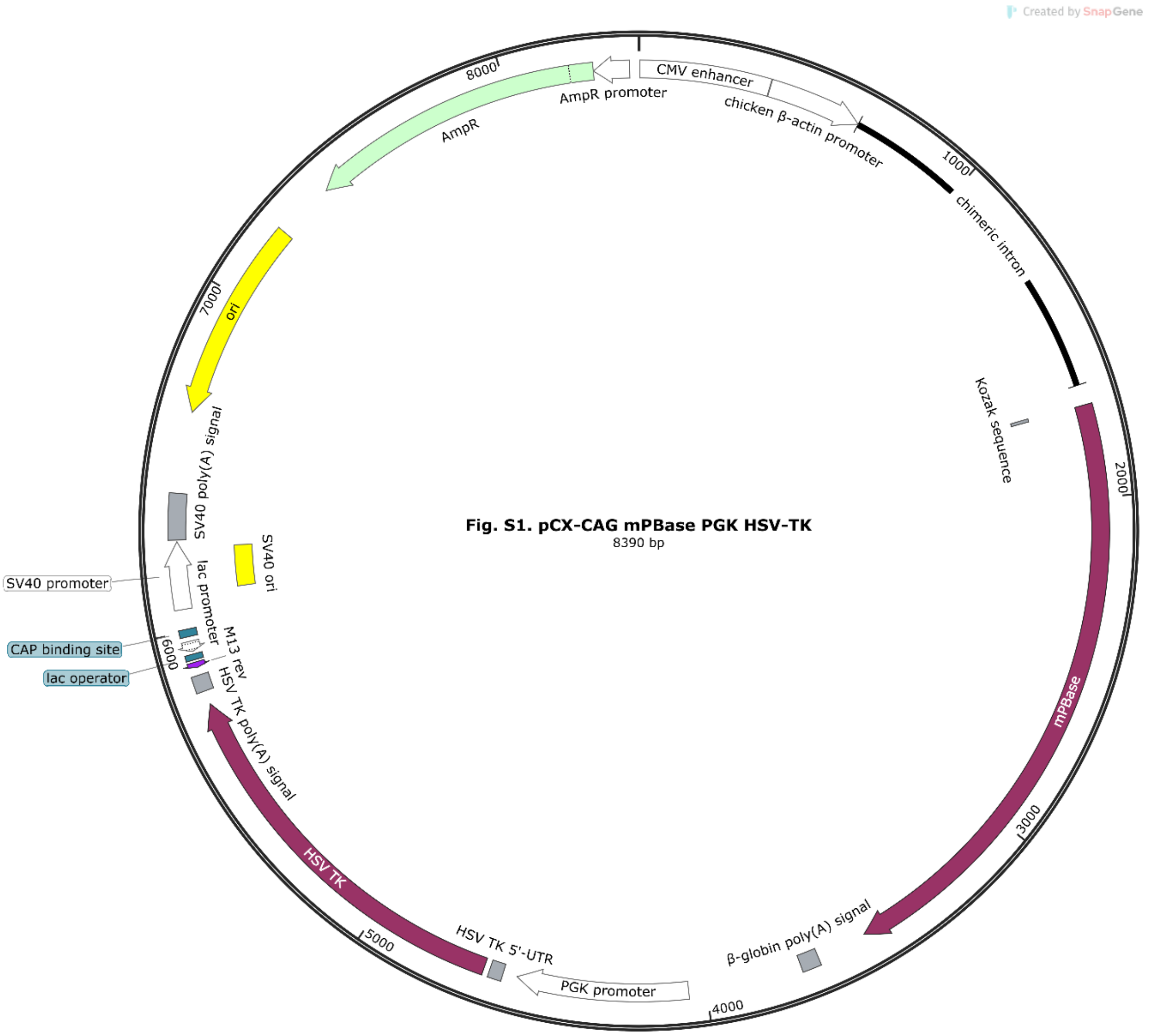


**Supplementary Figure S1. Vector map of *PiggyBac* transposase effector vector.**

Vector map of *PiggyBac* transposase (PBase) effector vector (pCX-CAG>mPBase-PGK>HSV-TK). The custom fragment containing a Kozak sequence attached to the coding sequence (CDS) of mammalian codon-optimised PBase (mPBase) was replaced from the EGFP of pCX-EGFP. The mPBase is regulated under the control of a CAG promoter and the HSV-TK is regulated under the control of a PGK promoter.

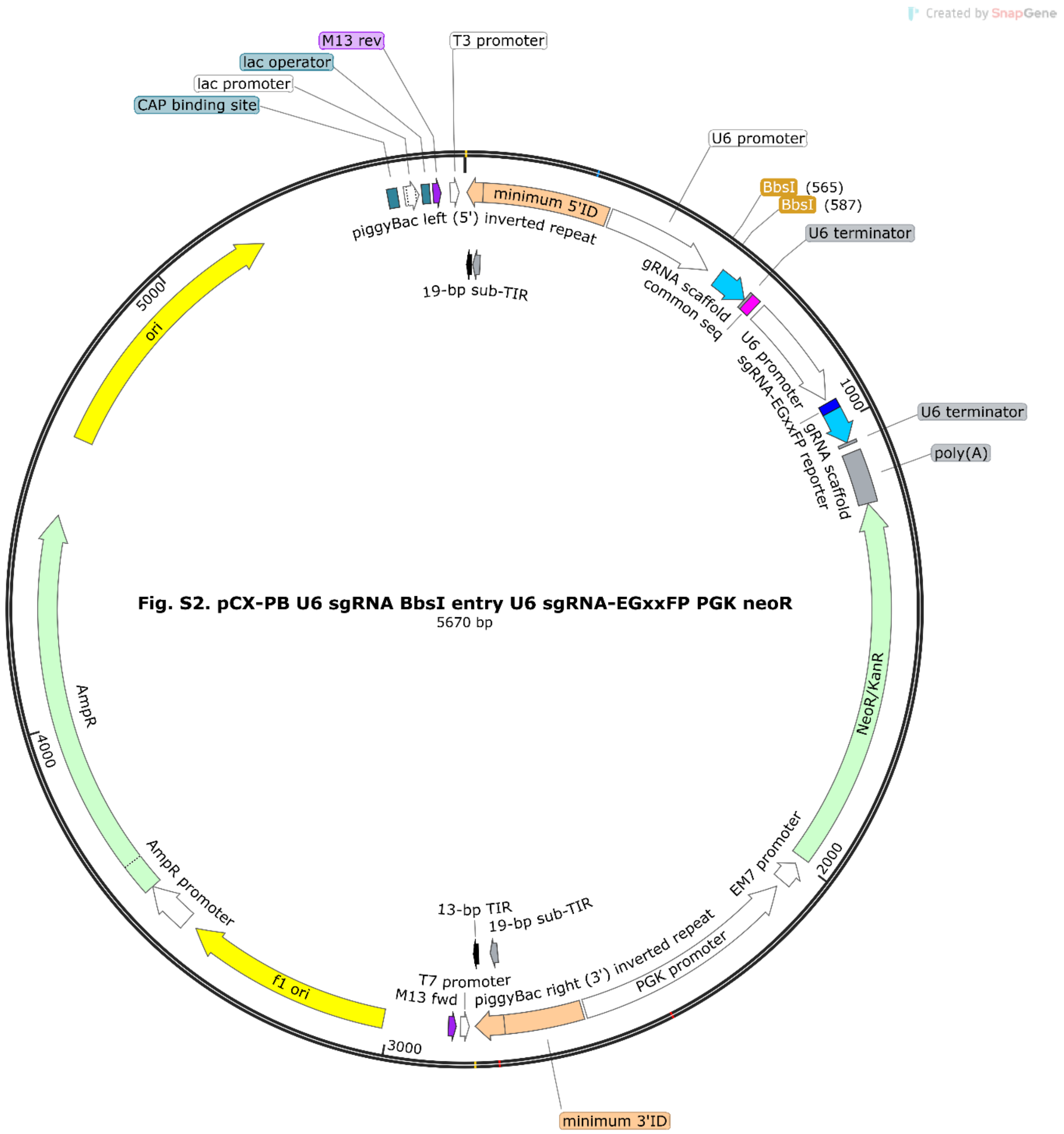


**Supplementary Figure S2. Vector map of PB single-guide RNA donor vector template.**

Vector map of PB single-guide RNA (sgRNA) donor vector template (pCX-PB U6>sgRNA-BbsI entry-U6>sgRNA-EGxxFP reporter-PGK>neo). Th PB-5’ inverted terminal repeat (ITR)-internal domain (ID) (PB-5’ITR-ID), the U6>sgRNA-BbsI entry containing BbsI sites that enables directional cloning of sgRNA oligos, U6>sgRNA-EGxxFP reporter, PGK>neo and PB-3’ITR-ID were replaced from the EGxxFP cassette of pCX-EGxxFP. The sgRNA-BbsI entry and sgRNA -EGxxFP reporter are regulated under the control of an U6 promoter and neo-resistance gene is regulated under the control of a PGK and an EM7 promoter.

Magenta box named common seq indicates reverse primer annealing site for targeted short next-generation sequencing (NGS).

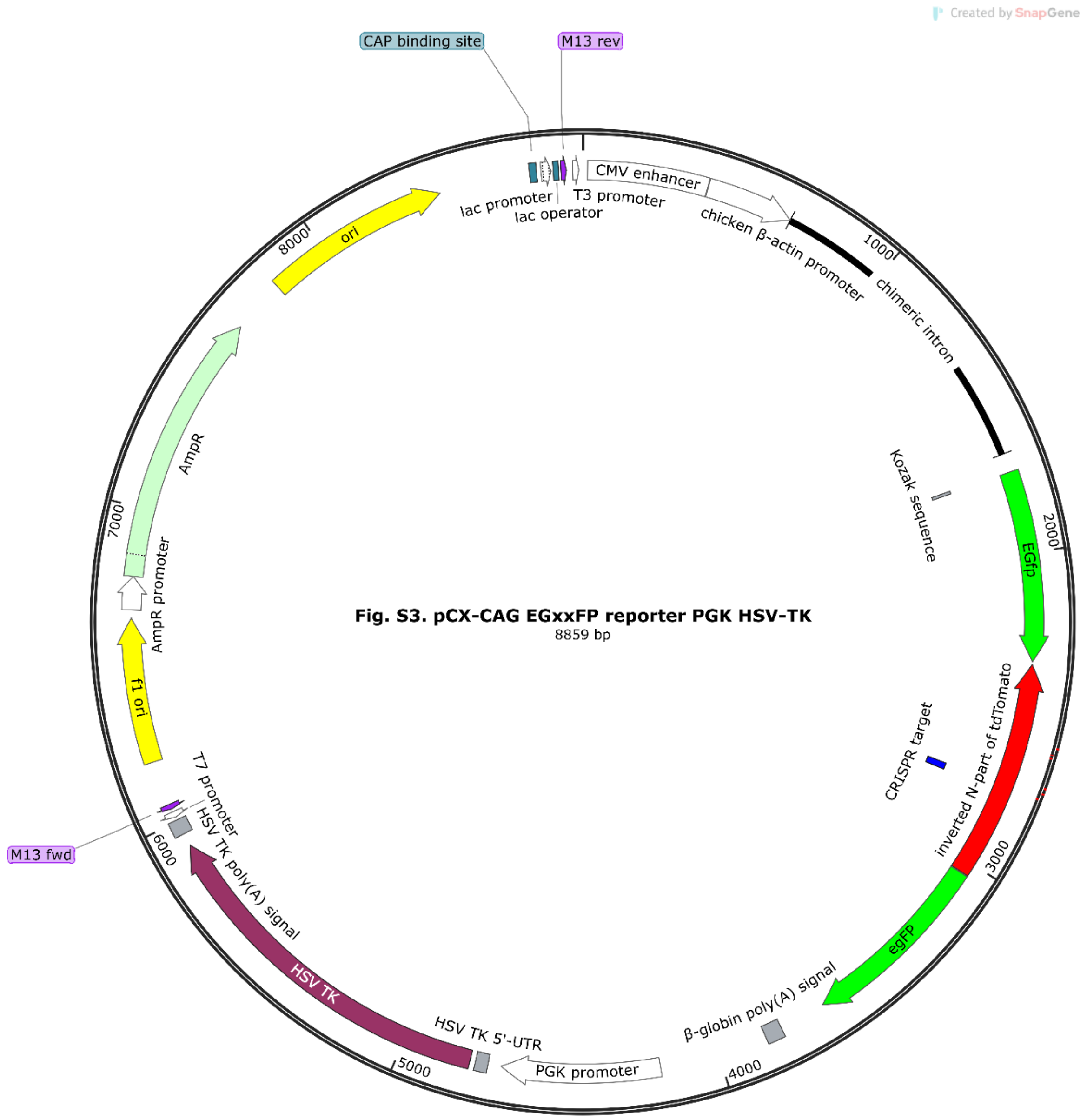


**Supplementary Figure S3. Vector map of EGxxFP reporter vector.**

Vector map of EGxxFP reporter vector (pCX-CAG>EGxxFP reporter-PGK>HSV-TK). The non-Chordata fragment (inverted N-part of tdTomato) containing the sgRNA-EGxxFP reporter target sequence was placed the N and C parts of the EGFP fragment of pCX-EGxxFP. The EGxxFP reporter is regulated under the control of a CAG promoter and HSV-TK is regulated under the control of a PGK promoter.

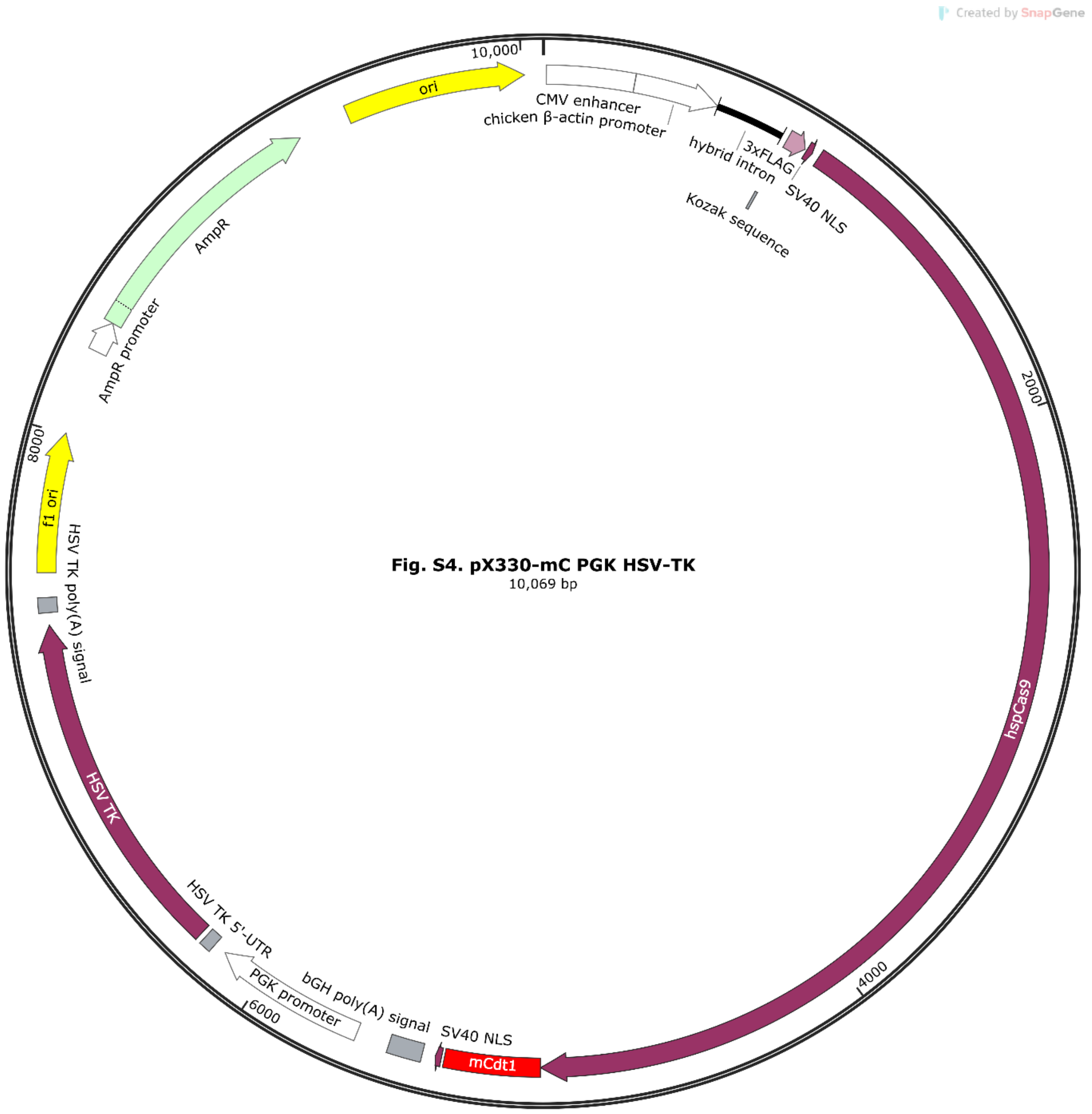


**Supplementary Figure S4. Vector map of Cas9 editor vector.**

Vector map of Cas9 editor vector (pX330-mC-PGK>HSV-TK). The human codon-optimised *Streptococcus pyogenes* Cas9 (hSpCas9)-Cdt1 (mouse) fusion protein is regulated by the control of a CBh promoter and HSV-TK is regulated under the control of a PGK promoter.

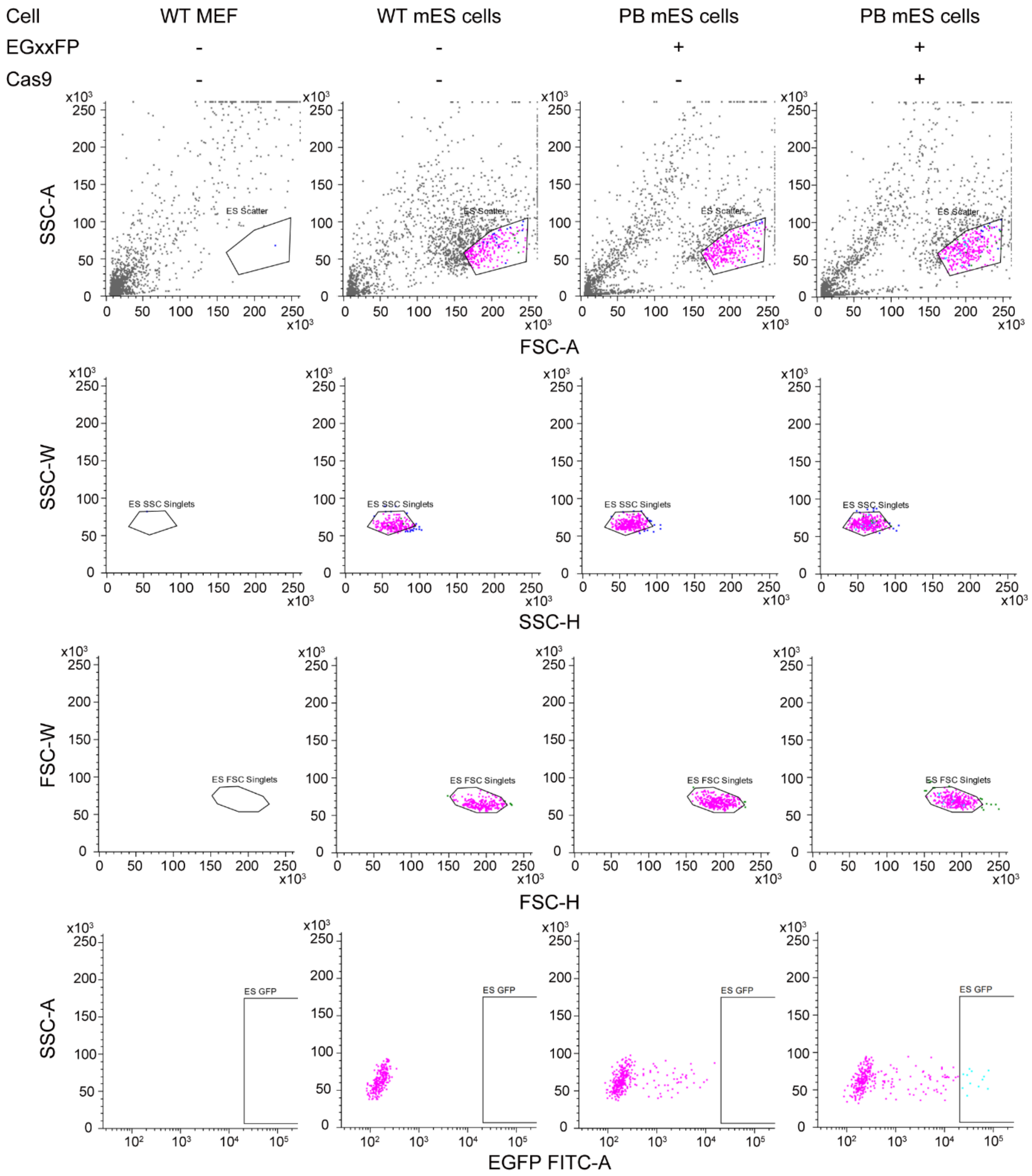


**Supplementary Figure S5. Gate setting for FACS.**

Represent gates setting for FACS (single-cell sorting). Upper table shows the conditions of transfection. Lower scatter graphs show total 2000 events in each transfection condition. The black frames in scatter graphs show the gates. To certainly sort only mES cells, the gates were set to capture no WT mouse embryonic fibroblast (MEF).
